# Dual Screen for Gut Metabolites Suppressing Enterobacterial Growth and Invasiveness Reveals Structure – Activity Relationships among Anti-Infective Indoles

**DOI:** 10.64898/2026.02.13.705707

**Authors:** Alexandra Bergholtz, Weifeng Lin, Amanpreet Kaur, Anjeela Bhetwal, Maria Letizia Di Martino, Jens Eriksson, Daniel Globisch, Mikael E. Sellin

## Abstract

The antibiotic resistance crisis makes characterization of new anti-infective molecules a pressing matter. Molecules that suppress bacterial growth and survival, virulence, or the combination of these traits, all warrant further exploration. Naturally occurring microbe-host ecosystems, such as the human gut, provide incompletely tapped resources in this regard. We developed a flexible platform to parallelly assess how gut metabolites affect the growth and epithelial cell invasion capacity of the enteropathogens *Salmonella enterica* Typhimurium (*Salmonella*) and *Shigella flexneri*. By screening a gut metabolite library, the assays identified multiple anti-infective compound classes and extended previously reported antibacterial activities for e.g. medium chain fatty acids, bile acids, purine nucleotides, and indole. Importantly, a targeted follow-up screen combined with chemical biology iterations showed how the anti-infective activity of indole is impacted by its derivatization. Specifically, a methyl group at either of the carbons of the indole scaffold potentiated the suppressive effect on type-III-secretion-system-mediated virulence, flagellar motility (for *Salmonella*), and growth, in a concentration-dependent manner. By contrast, N1-methylation markedly attenuated the activity of indole and its C-derivatized versions. The study, hence, offers assays for dual growth and virulence analysis of invasive enterobacteria exposed to anti-infective candidate molecules, and informs on structure–activity relationships among indole metabolites.

## Introduction

Pathogenic bacteria use their virulence factors to colonize host tissues, causing tissue destruction, inflammation, and potentially lethal systemic spread. Antibiotics remain our standard therapy for treating bacterial infections, but the spread of antimicrobial resistance mechanisms put current treatment options at risk (1). In the search for new chemical agents that selectively suppress or disarm pathogens, while keeping microbe-host ecosystems minimally altered, naturally occurring metabolites represent interesting starting points (2). Because of its staggering microbial and chemical complexity (3), the mammalian gut and its metabolome constitute a particularly relevant resource for such exploration.

Aggressive enterobacteria such as *Salmonella enterica* serovar Typhimurium (*S*.Tm) and *Shigella flexneri* (*S.fl*) colonize the gut lumen and invade intestinal epithelial cell (IECs) by means of Type-3-Secretion-Systems (T3SS) that work in concert with species- and strain-specific accessory virulence factors (4–7). Enterobacterial virulence is costly and therefore tightly regulated in response to external queues (8–13). The resident gut microbiota provides colonization resistance against incoming pathogens by competition for space and nutrients (14,15), stimulation of host counter defenses (16,17), or production of microbial warfare molecules (18,19). In addition, gut metabolites produced by either microbiota or host have been shown to modulate bacterial virulence (20). Short chain fatty acids (e.g. propionate and butyrate) can be present in high mM concentration in the gut and suppresses T3SS expression in *S.*Tm (21–23). Several medium- (MCFAs) and long-chain fatty acids (LCFAs) (24–28), bile acids (29,30), and the tryptophan-derived metabolite indole (31–34) have also been linked to suppression of either enterobacterial virulence or growth. Pathogen growth-suppressive activities have furthermore been recently assigned to e.g. adenine, adenosine (35) and their derivatives (36–38). These observations notwithstanding, the majority of the gut metabolome remains to be functionally mined for anti-infective properties.

Towards this goal, we present an imaging-based screening platform to assess the impact of diverse gut metabolites on *S*.Tm and *S.fl* growth and IEC invasion capacity. Combining this with a comprehensive and diverse library of singly-stocked gut metabolites revealed multiple metabolite classes capable of suppressing *S*.Tm invasiveness. A follow-up screen identified structural modifiers among indole derivatives, linking indole modification patterns to the anti-infective activity profile and potency. Most notably, we found that C-methylation at five positions (C2, C3, C5, C6, and C7) potentiates the suppressive effect of indole on both enterobacterial T3SS expression, flagellar motility, and growth. By contrast, N-methylation at position 1 drastically attenuated these indole activities. Taken together, this study offers a new strategy to resolve anti-growth versus anti-virulence activities of metabolites present in the gut and provides insights into indole structure–activity relationships.

## Results

### An assay system to quantify enterobacterial growth and IEC invasion following gut metabolite exposure

To screen gut metabolite effects on enterobacterial growth and virulence, we developed a 96-well-format assay to quantify *S*.Tm inoculum behavior and invasion of cultured epithelial cell layers (inspired by (39)). Human Caco-2 cells, or human enteroid-derived IECs, were seeded and matured into confluent monolayers. A *S*.Tm strain carrying the p*ssaG*-GFP reporter (turns GFP+ when arriving in the *Salmonella*-containing vacuole following invasion; (40,41)) was prepared and subcultured for 4h following a 1:20 dilution in the presence or absence of the tested metabolites. The subculture was used as inoculum to infect the epithelial monolayers (20min, MOI 50), followed by 3h incubation in gentamicin-containing medium to eliminate extracellular bacteria and allow p*ssaG*-GFP reporter maturation. Infected monolayers were counterstained followed by quantification of total IECs and intraepithelial *S*.Tm invasion foci (GFP+ spots) by a semi-automated microscopy pipeline (Fig 1A, Fig S1A-C; see Methods). OD600 of the 4h *S*.Tm subculture was measured in parallel to quantify effects on bacterial growth. For both scoring of *S*.Tm growth and invasion foci frequency, results were normalized to the corresponding data for unperturbed *S*.Tm *wt*/p*ssaG*-GFP, giving rise to the metrics “relative OD600” and “relative invasion” (Fig 1A; *wt* unperturbed = 1).

**Fig 1.**
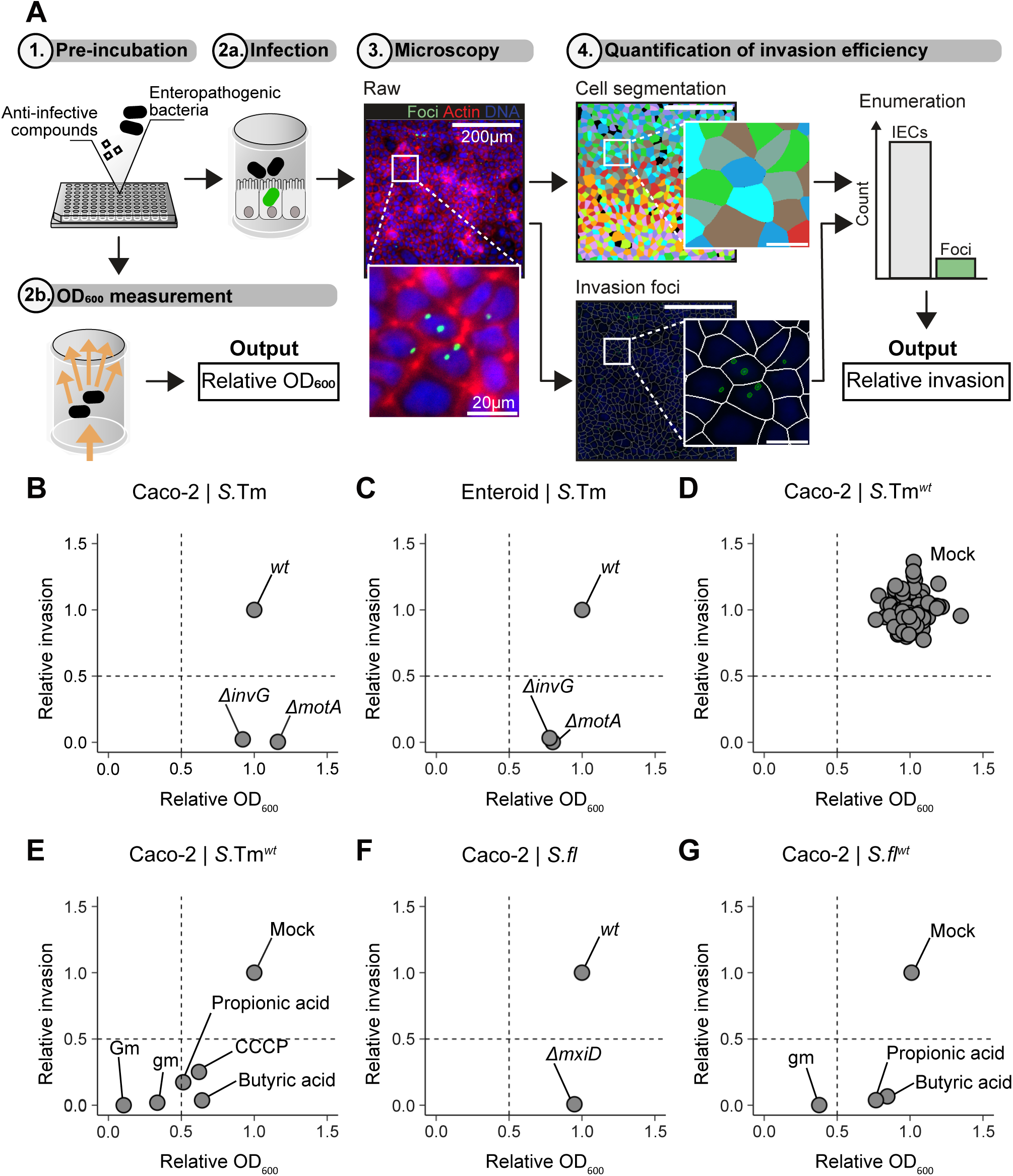
A 96-well format platform for dual quantification of enteropathogenic bacterial growth and epithelial cell invasion. **A.** Schematic illustration of the 96-well format IEC monolayer infection assay: (**1**) Enteropathogenic bacteria expressing the intracellular GFP reporter construct p*ssaG*-GFP (*S*.Tm) or p*UhpT*-GFP (*S.fl)* are incubated in presence or absence of metabolites/compounds of interest prior to **(2a)** infection of IECs and **(2b)** OD_600_ measurement. **(3)** The infected IEC monolayers are fixed in 2% PFA, stained with DAPI (blue) and Alexa Fluor 647 Phalloidin (red), and imaged using semi-automated fluorescence microscopy. **(4)** IECs and invasion foci are enumerated using the image analysis software CellProfiler in combination with the cell segmentation algorithm Cellpose. The obtained OD_600_ and invasion ratio for each infection are normalized to their corresponding *wt* or mock-treated control (“relative OD_600_” and “relative invasion”). See Methods for details. **B-G.** *S*.Tm or *S.fl* relative OD_600_ (x-axis) and relative invasion into IECs (y-axis). **B.** Caco-2 cell monolayers infected with T3SS-1 deficient *S.*Tm*^ΔinvG^*(n=28), non-motile *S*.Tm*^ΔmotA^* (n=32) and *S*.Tm*^wt^*(n=32). **C.** Undifferentiated human enteroid-derived monolayers infected with T3SS-1 deficient *S*.Tm*^ΔinvG^* (n=2), non-motile *S*.Tm*^ΔmotA^*(n=2) and *S*.Tm*^wt^* (n=3). **D.** Caco- 2 cell monolayers infected with *S*.Tm*^wt^* (n=91). Each dot represents the relative invasion ratio and relative OD_600_ from individual wells relative to the mean for all replicates. **E.** Caco-2 cell monolayers infected with *S*.Tm*^wt^* pre- incubated with anti-infective metabolites/compounds: butyric acid (10mM), propionic acid (10mM), CCCP (50µM), high dose gentamicin (Gm, 100µg/ml), low dose gentamicin (gm, 10µg/ml) (n=2), and mock-treated control (n=4). **F.** Caco-2 cells infected with T3SS deficient *S.fl^ΔmxiD^*(n=7) and *S.fl^wt^* (n=8). **G.** Caco-2 cells infected with *S.fl^wt^* pre-incubated with anti-infective metabolites/compounds: butyric acid (5mM), propionic acid (5mM), low dose gentamicin (gm, 10µg/ml) (n=2), and mock-treated control (n=3). Dotted lines depict the 0.5 cut-offs.

A set of control experiments were conducted to ensure robustness and resolution. The *S*.Tm T3SS-1 and flagella are well characterized virulence factors known to drive IEC invasion (4,42,43). In line with this, *S*.Tm*^ΔinvG^* (lacks a key T3SS-1 component; (44)) and *S*.Tm*^ΔmotA^* (incapable of flagellar movement; (45)) reporter strains exhibited normal relative OD600, but dramatically reduced relative invasion in Caco-2 monolayers (Fig 1B; invasion ratios of 0.0019 and 0.00031, respectively). Similar results were noted for human enteroid-derived IEC monolayer infections (Fig 1C). However, the total number of invasion events was markedly lower for any infection of the enteroid-derived IEC monolayers, particularly following their maturation in an enterocyte differentiation medium (Fig S1B-C; in line with our recent study; (46)). The low frequency of invasion events, combined with the propensity of non-transformed IECs to undergo swift cell death and extrusion upon *S*.Tm infection (41,45,47), makes the Caco-2 monolayers the more robust host cell model for the present application. Any screen comes with the risk of false positive results. To assess stochastic variability in the readouts, we carried out 91 parallel mock infection assays in Caco-2, with the *S*.Tm*^wt^* reporter strain pretreated only with solvent (0.5% DMSO in PBS), and compared to the average of all wells (Fig 1D). All 91 samples clustered around the mean (Fig 1D; mean±sd for relative OD600 0.957+/-0.092 and relative invasion 0.999+/-0.116). Based on this, conservative cut-offs of relative OD600 ≤0.5 and relative invasion ≤0.5 were chosen, which should yield less than 1% false positive results in a screening context. Applying those cut-offs, and from here using solvent (mock) treated *S*.Tm*^wt^*as reference, this assay system could readily pick up suppression of epithelial invasion by established gut metabolites and other chemical compounds (Fig 1E). Incubation with short-chain fatty acids (SCFAs; 10mM) that interfere with *S*.Tm T3SS-1 virulence (propionic acid and butyric acid; (21–23)), as well as Carbonyl cyanide *m*-chlorophenylhydrazone (CCCP; 50µM) that nullifies the proton-motive force for flagellar motility (48,49), reduced relative invasion (≤0.25). However, these only had modest effects on relative OD600, not reaching the cut-off (Fig 1E). Further as anticipated, gentamicin (10-100µg/ml) reduced both relative OD600 and relative invasion in a dose-dependent manner. This illustrates that the assay system has a low inherent noise and can reconfirm effects of known anti-infective metabolites and antibiotics on *S*.Tm growth and invasiveness.

Finally, we developed a similar assay for the related enterobacterium *S.fl*. The fluorescent reporter was exchanged for p*uhpT*-GFP (turns GFP+ upon bacterial arrival in the epithelial cell cytosol, the prevailing niche for *S.fl*;(50–52)). The assay also included a spin step after adding the *S.fl* subculture to the host cells, to compensate for *Shigella*’s lack of motility. Similar to for *S*.Tm, this assay system resolved differences in relative invasion between *S.fl^wt^*and *S.fl^ΔmxiD^* (lacks a key T3SS component) (Fig 1F). Moreover, short chain fatty acids again suppressed relative invasion, while a low dose of gentamicin decreased both relative OD600 and relative invasion (Fig 1G).

### Screening of a gut metabolite library for anti-infective activities

The *S*.Tm assay system was used to explore the anti-virulence and anti-growth properties of diverse human gut metabolites, implementing first an individually stocked library (MetaSci, SKU: MSIFEC0001) with 481 compounds present in human fecal samples. The library comprises of a variety of metabolites synthesized or modified by the gut microbiota, human biomolecules, and food breakdown products. The *S*.Tm*^wt^* reporter strain was pre-incubated with each metabolite at a 10mM concentration and the relative OD600 and relative invasion indexes assessed (as in Fig 1A), with mock-treated *S*.Tm*^wt^*as reference. Some metabolites exhibited signs of precipitation, predicted to cause an effective concentration <10mM in those pre-incubation cultures. Nevertheless, applying the ≤0.5 cut-offs, 30.4% of all metabolites tested were in this initial screen found to suppress the bacterium’s relative invasiveness and/or growth (Fig 2A-B). Out of these, 95 (65.1% of “hits”) suppressed invasiveness (lower right quadrant in Fig 2A; relative OD600 ≥0.5, relative invasion ≤0.5), whereas 51 (34.9% or hits) suppressed both parameters (lower left quadrant in Fig 2A; relative OD600 ≤0.5, relative invasion ≤0.5). Notably, not a single metabolite exclusively affected growth (upper left quadrant in Fig 2A). Any metabolite suppressing *S*.Tm growth during pre-incubation would be expected to yield fewer input bacteria in the IEC invasion assay and hence a correspondingly lowered invasion index. These results demonstrate the robustness and limited noise of our assays.

**Fig 2.**
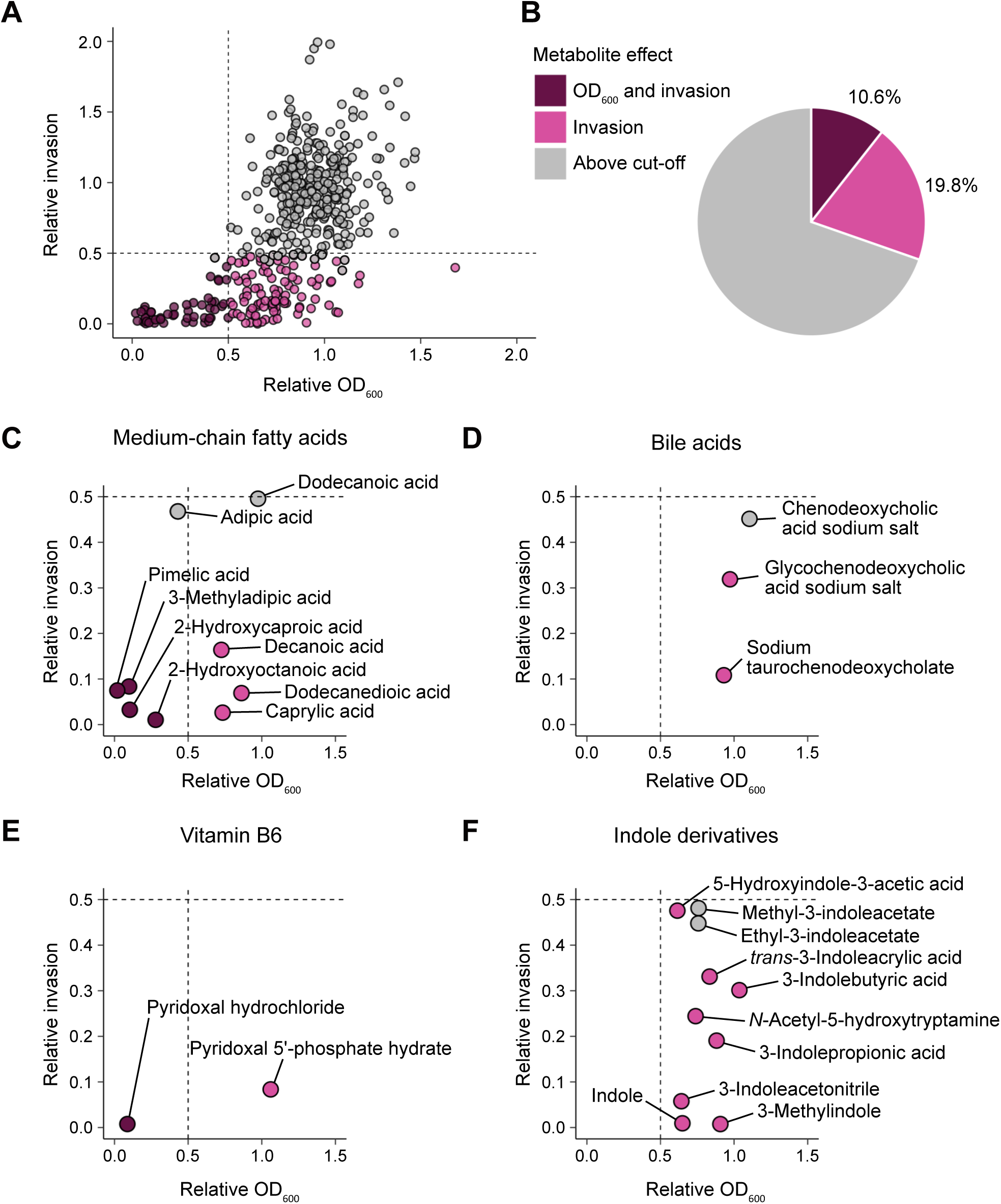
Screening of a gut metabolite library reveals functional classes and indole derivatives with variable anti- infective activity against S.Tm. **A.** *S*.Tm relative OD_600_ and relative invasion into IEC (Caco-2) monolayers after individual 4h pre-incubation with 481 metabolites from the Metasci human gut metabolite library. Each dot represents one metabolite result at 10mM (n=2 per condition). The fraction of metabolites with two replicates suppressing *S*.Tm relative invasion <0.5 is classified as hits. Relative OD_600_ <0.5 is used as cut-off for anti-growth activity. **B.** Categorization of metabolites from the library screen based on effect. Results are reported as percentage of the total number of screened metabolites. **C-F.** *S*.Tm relative OD_600_ and relative invasion into IEC monolayers after 4h pre-treatment with anti-infective metabolites from the library: **C.** medium-chain fatty acids, **D.** bile acids, **E.** vitamin B6, and **F.** indole derivatives (n=2 for each condition). Dotted lines depict the 0.5 cut-offs of relative OD_600_ (x-axis) and relative invasion (y-axis).

The final activity list for the 146 “hit” metabolites was mined for common themes and functional compound classes. This resulted in the re-discovery of specific metabolites that have previously been linked to bacterial virulence suppression (e.g. caprylic acid (53,54) and chenodeoxycholic acid (55)) (Fig 2C-D). Multiple medium-chain fatty acids (Fig 2C), and bile acid derivatives (Fig 2D), were among metabolites that reduced *S*.Tm invasiveness, with four medium-chain fatty acids also impacting *S*.Tm growth at the present concentration (Fig 2C). Vitamin B6/Pyroxidal metabolites were also identified (Fig 2E), as well as purine nucleotides (Fig S2A). Moreover, 19 metabolites caused >1.5-fold higher relative invasion (Fig S2B), signifying a boosting effect on *S*.Tm virulence.

Among metabolites suppressing *S*.Tm relative invasiveness below the ≤0.5 cut-off, no less than 10 indole derivatives were identified (Fig 2F). Indole itself is produced by gut microbiota members using tryptophan as precursor, and has been assigned suppressive effects on bacterial virulence (31–34,56). Intriguingly, however, our results revealed a highly variable activity profile of indole derivatives with different side groups. For example, although both indole and short-chain fatty acids on their own suppress *S*.Tm virulence, the coupling of either a propionate, or a butyrate side group to position 3 of indole resulted in weaker activity than for unmodified indole itself (Fig 2F). By contrast, 3-methylindole appeared highly potent. At the 10mM screening concentration, both indole and 3-methylindole essentially abolished *S*.Tm invasiveness, with partial effects also on growth (Fig 2F). Lowering the concentrations resulted in dose-dependent partial suppression of *S*.Tm invasiveness and hinted to a higher potency for 3-methylindole than for indole (Fig S2C-D). These findings suggest the suitability of the present assays to resolve structure–activity relationships among indole derivatives.

### A sub-screen of indoles provides structure–function insights and identifies a methylation position as a key anti-virulence determinant

To explore substituents of the indole scaffold in relation to the observed bioactivity, we conducted an additional sub-screen covering 40 indole derivatives. These included both metabolites covered in the original screen and other naturally occurring, as well as synthetic, indole derivatives with functional side groups at various positions of the bicyclic indole scaffold (Fig 3A). Dose-response curves (starting at 2mM) were generated for each compound and the concentration required to suppress *S*.Tm IEC invasion by 50% (IC50) calculated. The compounds were ranked according to their IC50 relative to unmodified indole, which under these conditions exhibited an IC50 of 681.4 µM when averaged over 3 replicate experiments (Fig 3B-D). In total 14 derivatives demonstrated higher potency (lower IC50), whereas nine showed lower potency (higher IC50) than unmodified indole (Fig 3A). The remaining 16 compound were even less potent and did not permit reliable IC50 quantification (Fig 3A, an inhibitory effect beyond 50% for the two highest concentrations was not reached, hence prohibiting IC50 determination). Corroborating the original screening results and observations by others (32), we found that bulky substituents such as e.g. addition of a butyrate or a propionate moiety in position 3, lowered the indole potency (Fig 3A).

**Fig 3.**
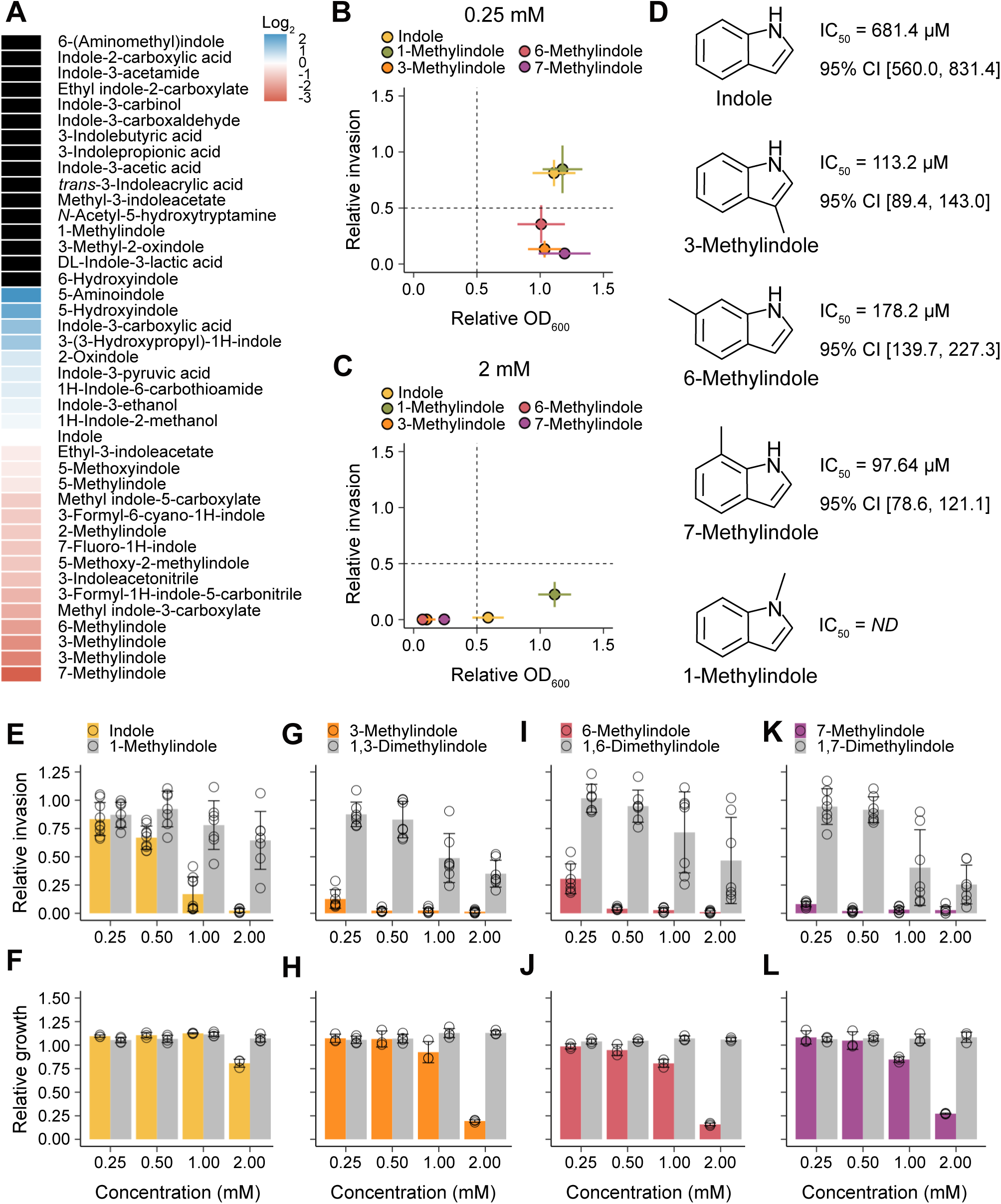
C-methylation of indole increases its anti-infective activity against *S*.Tm while N-methylation attenuates it. **A.** Heatmap of absolute inhibitory concentration required for 50% suppression (IC_50_) of *S*.Tm invasion into IEC (Caco-2) monolayers for the indicated indole-derivatives reported as log_2_ relative to indole (n=2). Dose-response curves for indole derivatives with ≤2 concentrations below the 50% predicted response are excluded from the IC_50_ determination (black box). **B-C.** *S*.Tm relative OD_600_ and relative invasion into IEC monolayers after 4h pre- treatment with indole (n=12), 1-methylindole, 3-methylindole, 6-methylindole or 7-methylindole (n=8) at the concentration of **B.** 0.25mM and **C.** 2mM. Dotted lines depict the 0.5 cut-offs. **D.** Chemical structure and IC_50_ with 95% confidence interval for indole (n=6), the top three most potent indole-derivatives; 3-methylindole, 6- methylindole and 7-methylindole (n=4). Chemical structure of 1-methylindole. ND, IC_50_ not determined (see Methods for rational). **E-L.** *S*.Tm relative invasion into IEC monolayers (n=7) and relative growth (area under the growth curve until end of exponential phase, n=3) after 4h pre-treatment with **E-F.** indole and 1-methylindole, **G-H.** 3-methylindole and 1,3-dimethylindole, **I-J.** 6-methylindole and 1,6-dimethylindole, or **K-L.** 7-methylindole and 1,7-dimethylindole. Data plotted represent mean±SD.

The hyperpotent indole compounds featured different side groups, but with a noticeable enrichment of methyl-indoles (Fig 3A). In addition to the previously identified 3-methylindole (included in duplicate from two different sources), also 2-, 5-, 6-, and 7-methylindole showed stronger suppression of *S*.Tm invasiveness than indole (Fig 3A-C). IC50 values were broadly similar across these derivatives, e.g. 113.2µM (95% CI: 89.4-143.0µM), 178.2µM (139.7-227.3µM), and 97.6µM (78.6-121.1µM) for 3-, 6-, and 7-methylindole, respectively (Fig 3D). By contrast, 1-methylindole (methylated at the amine of the pyrrole ring) was identified at the other end of the activity spectrum among the compounds that did not qualify for IC50 determination (Fig 3A-D). This suggests that C-methylation of the indole scaffold at essentially any position potentiates its suppressive effect on *S*.Tm invasiveness, while N-methylation at position 1 does the opposite.

Following on these findings, we examined if N1-methylation would reduce the effects also of the here identified hyperactive C-methylated indoles. We therefore synthesized three di-methylated indole derivatives by methylation of the N1-position of 3-, 6-, and 7-methylindole, respectively (see Methods for synthesis protocol). The potency of purified 1,3-dimethylindole, 1,6-dimethylindole, and 1,7-dimethylindole (purity validated in Fig S3A-C) to suppress *S*.Tm growth and IEC invasion was compared to the respective monomethylated compound. Synthetic N1-methylation of the indole scaffold itself abolished its suppressive effects (Fig 3E-F), in agreement with the results above (Fig 3A- D). Similarly, all three di-methylated indoles were dramatically attenuated compared to their 3-, 6-, or 7-mono-methylated counterparts (Fig 3G-L). Furthermore, indole and all C-mono-methylated derivatives suppressed *S*.Tm invasion also at concentrations where growth appeared unaffected (compare dose responses in Fig 3E/G/I/K with Fig 3F/H/J/L). These findings demonstrate that i) a methyl group at either of multiple C-positions increases indoles anti-infective activity, ii) both indole and C-methylated indole derivatives suppress *S*.Tm invasiveness at lower concentrations (<<1mM) and *S*.Tm growth at higher concentrations (>1mM), and iii) N-methylation at position 1 counteracts these activities across various indole derivatives.

### Methylation impacts indole derivative suppression of *Salmonella* virulence gene expression, motility, and enterobacterial invasivity

Indole has been ascribed anti-infective properties against both bacterial and fungal agents (34,57–61), including suppression of expression/functionality of virulence machinery such as T3SSs and flagella (31,32,61). To probe the mechanistic basis for our observations, quantitative RT-PCR was used to measure expression levels of representative *S*.Tm T3SS (*invG*, *sipA*) and flagellar (*fliC*, *fljB*) genes. In agreement with the phenotypic data (Fig 3), both 3-, 6- and 7-methylindole suppressed the expression of these virulence genes to a higher extent that indole itself. Expression of T3SS genes was overall affected at lower concentrations than flagellar genes for indole and all three C-methylated indole derivatives. Importantly, 1-methylindole again had a weakened effect, requiring higher concentrations for inhibition of T3SS expression, and causing essentially no inhibition at all of flagellar gene expression (Fig 4A-D). To formally evaluate how indole and the methylated derivatives affect flagellar motility, we monitored swimming dynamics in individual motile bacteria using differential interference contrast (DIC) time-lapse microscopy (62). The analysis confirmed that exposure to 1mM indole markedly reduced *S*.Tm median swimming speed (Fig 4E; 11.9µm/s) relative to mock-treated control (Fig 4E; 20.0µm/s). Again, 3-, 6- and 7-methylindole exhibited increased potency (Fig 4E; 8.3µm/s, 9.6µm/s and 9.0µm/s respectively), while no improved effect was observed for 1-methylindole (Fig 4E). This demonstrates that the methylation pattern is a key determinant of the potency of indole as an anti-*S*.Tm agent targeting multiple traits involved in both growth and virulence.

**Figure 4.**
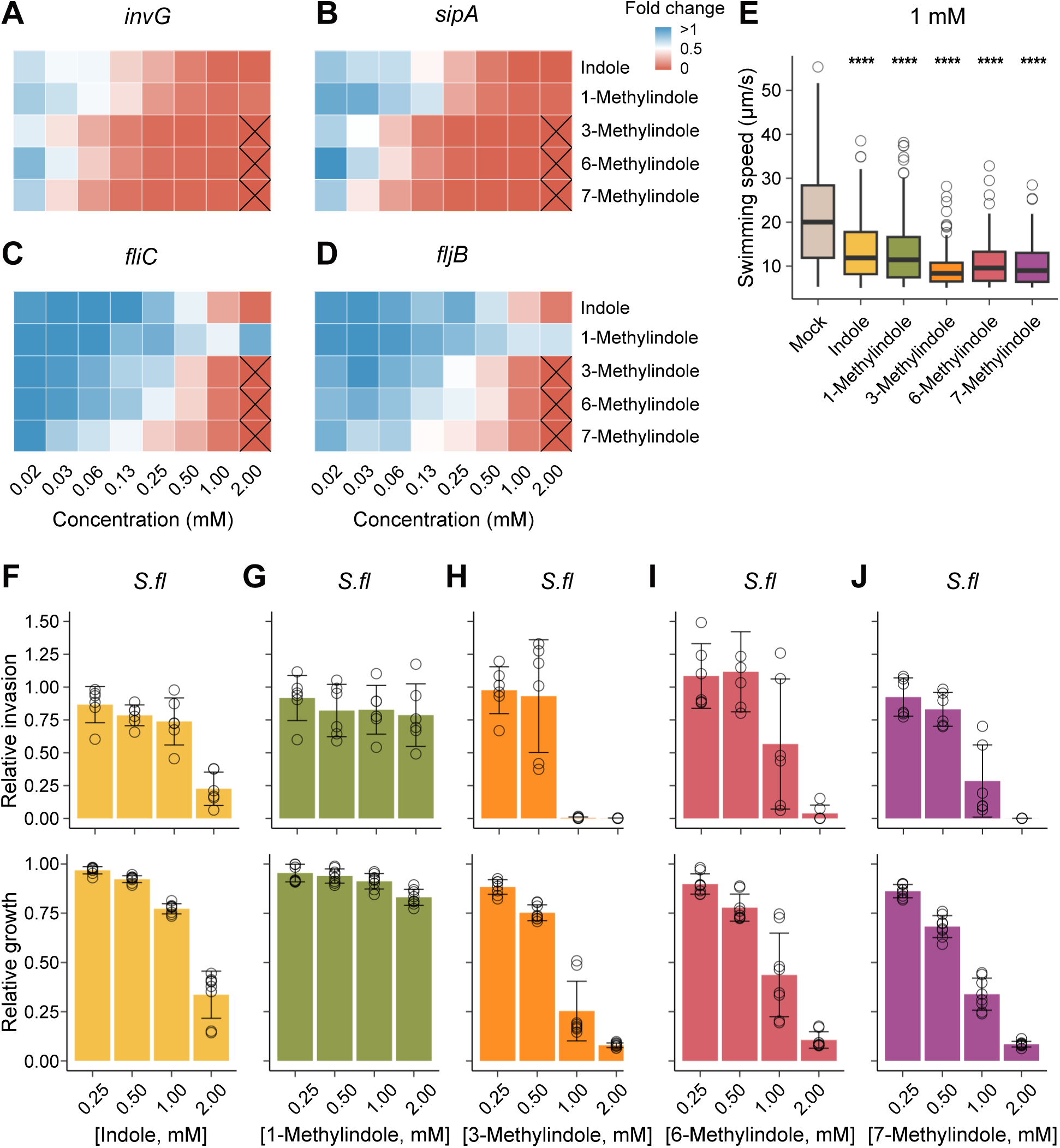
C-methylation of indole potentiates its suppressive effect on *S*.Tm virulence gene expression, flagellar swimming speed, and *S.fl* invasiveness. **A-D.** Heatmap of fold change in transcript levels of *S*.Tm virulence genes **A.** *invG*, **B.** *sipA*, **C.** *fliC* and **D.** *fljB*, after 4h treatment with indole, 1-methylindole, 3-methylindole, 6-methylindole or 7-methylindole relative to mock- treated control (n=3). Samples with strong growth-inhibitory effect were excluded from the analysis (“x”-marked box). **E.** Swimming speed atop glass of individual, motile *S*.Tm (defined as having a speed of >5µm/s, n≥90 for each condition) cultured in presence of 1mM of the corresponding indole derivative for 4h. The horizontal lines indicate the median and the enclosing boarders indicate 25^th^ and 75^th^ percentiles. The length of the whiskers indicate 1.5 times the interquartile range. Open circles represent outliers. Statistical analyses via Kruskal-Wallis test followed by Dunn’s test for multiple comparisons between mock- and indole-derivative-treated bacteria with Bonferroni comparison correction. ****p<0.0001. **F-J.** *S.fl* relative invasion into IEC (Caco-2) monolayers (n=6 per condition) and relative growth (area under the growth curve until end of exponential phase, n=8) after 2.5h pre-incubation with **F.** Indole, **G.** 1-methylindole, **H.** 3-methylindole, **I.** 6-methylindole or **J.** 7-methylindole.

Finally, to identify if the methylation-position-dependent augmentation/attenuation of the indole activity generalizes beyond *S*.Tm, we assessed the impact of the compound series on *S.fl* growth and invasivity, using the adapted assay system (see Fig 1A, F-G). It should be noted that in contrast to *S*.Tm, *S.fl* can (in the case of some strains) synthesize indole endogenously and also lacks flagellar motility (63,64). We found that the overall sensitivity to indole derivatives was lower for *S.fl* than for *S*.Tm (Fig 4F-J top panels, compare to Fig 3E, G, I, K). Nevertheless, both the relative IEC invasion and relative growth of *S.fl* was suppressed by indole at concentrations of ∼2mM (Fig 4F). Importantly 3-, 6- and 7-methylindole were yet again hyperpotent with detectable effects at 1mM (Fig 4H-J), while 1- methylated indole neither hampered *S.fl* invasivity, nor growth, at any of the 0.25-2mM concentrations (Fig 4G). We conclude that C-methylation of indole at a variety of positions broadly potentiates its anti-enterobacterial effects, whereas N1-methylation has the opposite outcome.

## Discussion

The rapid spread of antimicrobial resistance (AMR) poses one of this century’s leading threats to human and animal health (1). An improved understanding of molecules that target microbial growth or virulence is vital to better understand these processes and to counter the threat of AMR. Although several classes of microbiota-derived metabolites with anti-virulence activity have been identified (20), the gut metabolome remains an incompletely explored resource (65,66). Importantly, traditional methods for assessing antimicrobial activity, e.g. broth microdilution and disc diffusion assays, do not detect anti-virulence activities, which emphasizes the need for developing tools to dissect growth-suppressing versus virulence-suppressing activities of candidate molecules. To that end, we here established an imaging-based screening platform for parallel quantification of metabolite effects on bacterial growth and IEC invasion – a central virulence trait among Gram-negative enterobacteria.

Among the 481 gut metabolites initially screened for their anti-infective activity, our assays reconfirmed multiple classes of metabolites previously reported to influence *S*.Tm virulence, including MCFAs (54), LCFAs (24,25), bile acids (29,30,55), vitamin B6 (67) and purine nucleotides (68). The screen also re-identified indole (56), and an additional 9 indole derivatives, with variable anti-virulence potency against *S*.Tm. Extending on these findings, a sub screen of 40 indole derivatives revealed that the composition and position of side groups heavily impacted the anti-growth and anti-virulence properties of indole, hence expanding our understanding of structure-activity relationships within this molecule class (69–73). Notably, while a C-methyl modification (to e.g. positions 2, 3, 5, 6, 7) enhanced both the anti-virulence and anti-growth activity of indole against *S*.Tm and *S.fl*, addition of a methyl group to N1 rendered the molecule dramatically less potent in suppressing IEC invasion and completely eliminated its growth-inhibitory effect. The suggested importance of the N1-position chemical composition was further confirmed by N-methylation of the hyper-active C-methylated indoles, which in all cases led to loss of anti-infective activity. By contrast, a recent screen of indole derivatives against *C. parvum* demonstrated that parasite growth was inhibited by most derivatives regardless of substituent modification (73). However, that study investigated effects on a eukaryotic parasite in the infected host cell context, and did not include indoles substituted on the N1-position.

Indole has been shown to efficiently interact with and traverse membranes (74,75), and to act as a protonophore dissipating the proton motive force (PMF) by facilitating H+ flux across the cytoplasmic bacterial membrane (76,77). Since C-methylation will increase the overall lipophilicity, and since we found essentially identical effects of a C-methyl group at various positions, it appears likely that C-methylation drives accumulation and/or potentiates the membrane-directed activity of indole. It is further noteworthy that C-methylated indoles have a slightly lower predicted pKa than indole, which in addition could facilitate proton dissociation. Taken together, the potentiation of such activities to destabilize membranes and interfere with energy utilization, provides a plausible explanation to why C2-, 3-, 5-, 6-, and 7-methylation augments the suppressive effects of indole on *S*.Tm and *S.fl*. Conversely, we propose that N1-methylation, while also predicted to increase the overall lipophilicity, will make indole unable to act as a protonophore due to the disruption of the NH-group at this position.

Membrane damage and dissipation of the PMF can in a straight-forward way explain suppressive effects on bacterial growth and flagellar motility. However, at lower concentrations than those needed for *S*.Tm growth inhibition, indole and its C-methylated derivatives still suppressed virulence gene expression, particularly for SPI-1 genes. Moreover, even N-methylated indole (that had essentially no growth-inhibitory effect) was capable of suppressing SPI-1 gene expression at <1mM concentrations. These observations hint towards additional indole activities impinging on virulence gene expression and motility, which gains support in recent literature (31–33,78). To map and resolve the multiple potential molecular actions of indole derivatives on the deeper chemical level, and how they can be individually tweaked by either C- or N-position substitutions, constitutes exciting avenues for future research.

In summary, this study offers a platform for flexible dissection of anti-growth versus anti-virulence activities of metabolites and synthetic compounds that target enterobacteria. Moreover, our work provides foundational structure – activity insights that will facilitate the advancement of indole derivatives as research tools, or as scaffolds for future anti-infective development.

## Methods

### Bacterial strains, plasmids, and culture conditions

The bacterial strains and reporter plasmids used in this study are listed in Table S1. All *Salmonella enterica* serovar Typhimurium (*S.*Tm) strains were of SL1344 background (SB300, streptomycin resistant) (79). Besides the wild-type (*wt*), the previously characterized *ΔinvG* and *ΔmotA* mutant strains were used (44,45). All *Shigella flexneri* (*S.fl*) strains were of M90T background (80). Besides the *wt*, the previously described *ΔmxiD* mutant strain was used (81). For infections in the 96-well plate format, the indicated strain carried either the intracellular pM975v (p*ssaG*-GFPmut2) or pZ1400 (p*uhpT*-GFP) reporter plasmid (40,41,50,51). *S*.Tm strains were grown at 37°C for 12h in Luria Bertani broth (LB) with 0.3M NaCl (Sigma-Aldrich) supplemented with appropriate antibiotics on a rotating wheel (20rpm). A 1:20 dilution was subcultured in fresh LB 0.3M NaCl broth with- or without the indicated metabolites or anti-infective compounds, and grown in 96-well plates (Sarstedt #83.3924) at 37°C for 4h on a rotating table (40rpm). *S.fl* strains were grown at 30°C for 16h in LB with appropriate antibiotics and subcultured 1:50 in fresh LB with- or without metabolites or anti-infective compounds, and grown in 96-well plates at 30°C for 15min followed by 2h 15min at 37°C on a rotating table (40rpm). The total volume of the subculture in the 96-well plates was 100µl. For inoculum preparation, the 4h *S*.Tm subculture was diluted to 1.0x10^8^ CFU per ml in DMEM GlutaMAX (Gibco) with 10% heat inactivated FBS (Gibco) and 0.1mM non-essential amino acids (NEAA) (Gibco) (for Caco- 2 infections) or DMEM/F12 (for enteroid infections), of which 50µl was used for each infection, resulting in 5x10^6^ CFUs per IEC monolayer. The 2h *S.fl* subculture was diluted to 2.5x10^7^ CFU per ml in DMEM GlutaMAX with 10% heat inactivated FBS (Gibco) and 0.1mM non-essential amino acids (NEAA) (Gibco) of which 20µl was used for infection, resulting in 5x10^5^ CFUs per IEC monolayer.

### Ethics statement

Enteroids were established in prior studies (45,82), using human jejunal tissue resected during bariatric surgery. Enteroid culture and experimentation procedures were approved by the relevant governing body (Etikprövningsmyndigheten, Sweden) under license numbers 2010-157 (addenda 2010-157-1 and 2020-05754) and 2023-01524-01. Data presented in this study derive from enteroid culture Hu18-9jej.

### Caco-2 cell culture and monolayer establishment

Caco-2 cells were cultured in DMEM GlutaMAX (Gibco) with 10% heat inactivated FBS (Gibco), 0.1mM non-essential amino acids (NEAA) (Gibco) and 1x PenStrep (Gibco), at 37°C and 10% CO2. Caco-2 cell cultures were passaged every 2-3 days at a ratio of 1:4-1:8. Briefly, cells were detached using Trypsin-EDTA (Gibco) for 10min at 37°C and then spun down at 250xg for 4min. The cell pellet was resuspended in fresh culture medium and distributed in T-75 flasks. For infections in the 96-well plate format, trypsinized, resuspended Caco-2 cells were enumerated in a Bürker chamber and seeded at 1.43x10^5^ cells/cm^2^ (confluent monolayers, *S*.Tm infection) or 1.79 x10^4^ cells/cm^2^ (sub-confluent monolayers, *S.fl* infection) in a black 96-well culture plate (ibidi #89626) 2 days prior to infection. The medium was refreshed one day post seeding/prior to infection and antibiotics were omitted from the medium on the day of infection.

### Human enteroid culture and monolayer establishment

Human jejunal enteroids and culture conditions used in the study were characterized extensively in previous studies (45,82,83). Enteroids were cultured in 75% Matrigel (Corning) domes overlaid with human organoid growth medium (hOGM) (StemCell) with 1x PenStrep (Gibco) and incubated at 37°C and 5% CO2. Enteroid cultures were passaged every 6-8 days at a ratio of 1:8. Washing was performed in DMEM/F12/0.25% BSA and included centrifugation at 300xg for 5min at 4°C. For splitting, enteroids were manually disrupted by pipetting in gentle cell dissociation reagent (StemCell), followed by washing. Fragments were resuspended in hOGM and Matrigel at a ratio of 1:3 and divided into 50µl domes in 24-well plates. The overlying hOGM medium was exchanged every 2-4 days. For infections in the 96-well plate format, enteroids were dissociated into single cells using TrypLE Express (Gibco) for 5-10min at 37°C and then spun down at 300xg for 5min at 4°C. The cell pellet was washed one time and subsequently resuspended in hOGM with 1x PenStrep and 10µM Y-27632 (ROCK inhibitor, StemCell). Cells were enumerated in a Bürker chamber and seeded at 4.46x10^5^ cells/cm^2^ in a black 96- well culture plate (ibidi #89626). The ROCK inhibitor was omitted from the medium 2 days post seeding. For an undifferentiated stem cell-like phenotype, monolayers were maintained in hOGM with 1x PenStrep until infection on day 3 post seeding. For differentiation into an enterocyte-like phenotype, the hOGM medium was replaced to human enteroid differentiation medium (hODM) (StemCell) with 1x PenStrep on day 3 post seeding and maintained until infection on day 7. The medium was replaced with DMEM/F12 without antibiotics on the day of infection.

### Chemical synthesis

All reagents and solvents were purchased from Sigma-Aldrich or Fischer Scientific and were used without further purification. Solutions were concentrated *in vacuo* on a Heidolph rotaru evaporator. Thin Layer Chromatography (TLC) was performed on silica gel 60 F-254 plates. Visualization of the developed chromatogram was performed using fluorescence quenching. Chromatographic purification of products was accomplished using flash column chromatography on Merck silica gel 60 (40−63 μm). NMR spectra were recorded on Agilent 400 MHz spectrometer (1H NMR: 400 MHz, 13C NMR: 100 MHz). Chemical shifts are reported in parts per million (ppm) on the δ scale from an internal standard. Multiplicities are abbreviated as follows: s = singlet, d = doublet, t = triplet, q = quartet, m = multiplet. High-resolution mass spectra were acquired in positive mode at a mass range of *m/z* = 50– 1200. NMR spectra and further chemical synthesis details can be found in the Supplemental material.

### Platform for quantification of epithelial invasion by enteropathogenic bacteria in 96-well format

The IEC monolayers grown in black 96-well culture plates were washed and incubated in 150 µl of the indicated medium without antibiotics for 2h prior to infection. Inoculum of the indicated reporter *S*.Tm strain was prepared from the 4h subculture as described above, and added atop the IEC monolayers followed by incubation for 20min at 37°C with 5% CO2 (enteroids) or 10% CO2 (Caco-2 cells). Inoculum of *S.fl* was prepared from the 2h subculture as described above, and added atop the Caco-2 monolayers. The plate was centrifuged at 700xg for 10min prior to incubation for 45min at 37°C with 10% CO2. At the time of infection, the bacterial subculture optical density at 600 nm (OD600) was measured using a Tecan infinite M200 microplate reader. After the infection, the IEC monolayers were washed twice in DMEM/GlutaMAX (Caco-2 cells) or DMEM/F12 (enteroids) and incubated in the indicated medium supplemented with 200µg/ml gentamicin until 3h post infection at 37°C with 5% or 10% CO2. The IEC monolayers were then fixed in 2% paraformaldehyde for 20min, washed twice with DPBS (Gibco), and stained with DAPI (1µg/ml, Sigma Aldrich) and phalloidin-AlexaFluor647 (1U/ml, Molecular probes) in 0.1% Triton X-100 for 30min. After staining, the wells were imaged on a Nikon Ti2 microscope equipped with a 40x/0.6 NA Plan Apo air objective (pixel size 0.29µm) and a back-lit sCMOS camera (Prime 95B, Photometrics). Fluorescence was excited using a Spectra-X light engine (Lumencor), DAPI (390/22x), GFP (475/34x), Alexa647 (ET640/30x). A quadruple bandpass dichroic and emission filter (Chroma 89402bs & 89402m) was used for DAPI, GFP, and Alexa647 detection. Nine images per well in a 3x3 grid were automatically acquired using the µManager-high content screening plugin HCS Site generator (84). Images were analyzed with the open-source software CellProfiler (85) in combination with the segmentation algorithm Cellpose (86). Briefly, flatfield correction images were generated according to previous work by Model and Burkhardt (87). IECs segmentation was performed on Alexa647 and DAPI intensity images using Cellpose (v.1.0 and v.2.0). Invasion foci were detected using manually determined threshold values for pixel size and fluorescence intensity in the GFP channel with the IdentifyPrimary module. The threshold for fluorescence intensity was determined based on the background noise from non-infected control wells. GFP fluorescence signal outside of the segmented IECs were removed using the MaskObject module. The invasion ratio was calculated as the ratio between the number of invasion foci and the number of IECs per well. The relative invasion was calculated as the invasion ratio for the indicated strain or treatment relative to *wt* or mock-treated control. The background OD600 value from the LB-medium was subtracted from the subculture OD600. The relative OD600 was calculated as the OD600 for the indicated strain or treatment relative to *wt* or mock-treated control.

### Human gut metabolite library screen

Each compound from the human gut metabolite library (MetaSci Inc. Ontario, SKU: MSIFEC0001) was dissolved at 100mM in DPBS (Gibco) supplemented with 5% DMSO and stored at -20°C overnight. On the day of the experiment, the metabolite solutions were thawed for 30min in room temperature, followed by centrifugation at 14’000rpm for 1.5min and thereafter added to the *S*.Tm subculture as described above (final DMSO concentration 0.5%). After 4h of growth at 37°C on a rotating table (40rpm), the pretreated *S*.Tm inoculums were used to infect Caco-2 cell monolayers in the 96-well plate infection assay as described above. Reference metabolites were initially screened across a range of concentrations, after which 10mM was selected as the standard concentration for the main screen, with 1mM concentration samples also included for internal reference. All screening experiments were performed in duplicates. Each plate assayed 22 different metabolites from the library, four mock-treated controls, two gentamicin-pretreated controls (100µg/ml) and two uninfected control wells. Metabolites with two replicates suppressing *S*.Tm relative invasion <0.5 were classified as hits. Relative OD600 <0.5 was used as cut-off for anti-growth activity. A selection of hit metabolites were diluted in a 7-point 1:2 serial dilution series starting from the highest concentration used in the main screen and validated in the 96-well plate infection assay as described above.

### Indole derivative screen

Each indole derivative (Sigma-Aldrich Sweden) was dissolved at 2.1mM in LB 0.3M NaCl supplemented with 0.5% DMSO and thereafter centrifuged at 14’000rpm for 5min on the day of the experiment. LB 0.3M NaCl/0.5% DMSO was used for mock-treated control. *S*.Tm subcultures were prepared in a 96- well plate format as described above with- or without the indicated indole derivative and incubated at 37°C for 4h on a rotating table (40 rpm) (final DMSO concentration 0.475%). The pretreated *S*.Tm inoculums were used to infect Caco-2 cell monolayers in the platform for quantification of IEC invasion as described above. All indole derivatives were screened in 8-point 1:2 serial dilution series starting from 2mM and in duplicates. Each plate assayed four different indole-derivatives, one internal dilution series of indole, 12 mock-treated controls, two gentamicin-pretreated controls (100µg/ml) and two uninfected control wells.

### Indole IC50 determination

The absolute inhibitory concentration of indole and indole derivatives required for 50% suppression of *S*.Tm invasion into Caco-2 cells (IC50) was calculated from dose-response curves generated from the invasion ratio data obtained in the indole derivative screen. The 100% response was defined as the mean invasion ratio for mock-treated controls and the lowest concentration of indole-derivatives within the same experiment. The 0% response was defined as the mean invasion ratio obtained from high dose gentamicin-treated controls (100µg/ml). A non-linear curve fit was applied to the dose-response curves using GraphPad Prism (settings: log(inhibitor) vs normalized response, standard slope: Hillslope = -1). Data presented as IC50 relative to the internal indole-treated control. Indole derivatives with <2 concentrations below the predicted 50% response were excluded from the IC50 determination.

### Enterobacterial growth assay

*S*.Tm or *S.fl* subcultures were prepared in a 96-well plate format as described in the indole derivative screen. Measurements of the culture’s OD600 were recorded automatically every 15min during 5h 45min (growth until late exponential phase) in a Tecan Spark 10M Multimode Plate Reader. The 96- well plates were kept with closed lids, in a moist chamber, at 37°C and with continuous shaking between the measurements. The background OD600 values from the medium were subtracted from the bacterial culture OD600. Area under the growth curve (AUC) was calculated in GraphPad Prism (10.4.0) (settings: baseline = 0). Data presented as AUC relative to mock-treated control.

### RNA extraction

*S*.Tm subcultures were grown in a 96-well plate format as described in the indole derivative screen. The 4h subcultures from three wells were pooled together, spun down and resuspended in TES buffer (10mM Tris HCl pH8, 1mM EDTA, 150mM NaCl). Total RNA was extracted from three independent overnight cultures using acid phenol (Sigma Aldrich). All centrifugation steps were performed at 14’000rpm at 4°C. Briefly, the samples were incubated with 10% SDS at 95°C for 5min and thereafter cooled on ice. Samples were incubated at 65°C for 3min followed by addition of 200µl acid phenol. Samples were incubated at 65°C for additional 10min and vortexed every 2min. Samples were centrifuged for 10min and the aqueous phase transferred to a new tube and mixed with 180µl chloroform. Samples were incubated at 65°C for 5min followed by 5min centrifugation. The aqueous phase was then transferred to a new tube before addition of 375µl ice-cold 100% EtOH. Samples were incubated at -80°C for at least 40min and thereafter centrifuged for 30min. Pellets were washed with ice-cold 70% EtOH, centrifuged for 10min, air dried and dissolved in sterile water. The extracted RNA concentration was measured with Nanodrop and the RNA quality was checked by agarose gel electrophoresis. Total RNA was DNaseI (Thermo Fischer) treated at 37°C for 30min. Samples with strong growth-inhibitory effect were excluded from the analysis.

### Real-Time quantitative PCR

Oligonucleotide primers used in this study are listed in Table S2. Total RNA was reverse transcribed with high-capacity cDNA RT kit (Thermo Fischer) as described previously (12,88). Real-Time quantitative PCR was performed with Maxima SYBR Green (2X) (Thermo Fischer). Reactions were run on a CFX384 Real-Time Systems Instrument (BioRad). The data was analyzed with the Bio-Rad CFX Maestro Software. Transcript levels were calculated using the 2^-ΔΔCT^ method (89). The housekeeping gene *recA* was used for normalization. Data presented as transcript levels relative to mock-treated control.

### Analysis of single-bacterium swimming speed

*S*.Tm subcultures were grown in presence or absence of 1mM of the indicated indole-derivative in a 96-well plate format as described in the indole derivative screen. The 4h subcultures were diluted 1:250 in fresh medium containing the corresponding indole-derivative and subsequently added to a black-walled glass bottom 96-well plate (CellVis #P96-1.5H-N). Live DIC imaging was initiated 10min after addition to the plate, on a Nikon Ti2 microscope using the 60x/0.7 S Plan Fluor air objective (pixel size 183nm). Images were acquired at 250ms intervals for 6 frames (1.25s). The time-lapse series were imported to ImageJ and corrected for uneven background before single-bacterium swimming speed was tracked using the TrackMate plugin (90) of ImageJ as previously described (62). Tracks with swimming speed <5µm/s (defined as non-motile) were excluded from the analysis.

## Acknowledgements

The authors are grateful to members of the Sellin and Globisch laboratories, and to Professor Per Artursson (Department of Pharmacy), for fruitful discussions. This work was funded by the Swedish Foundation for Strategic Research (FFL18-0165 to MES), the Swedish Research Council (2022-01590 to MES, 2020-04707 to DG), the Swedish Cancer Society (25 4898 Pj to DG), Science for Life Laboratory (SciLifeLab Fellows program grants to DG and MES), and an Uppsala Antibiotic Center PhD student fellowship (to ABe, MES).

## Contributions

Conceptualization: ABe, DG, MES; Methodology: ABe, WL, AK, ABh, MLDM, JE; Investigation: ABe, WL, AK, ABh; Formal analysis: ABe, WL, JE, ABh; Interpretation: ABe, DG, MES; Resources: DG, MES; Project administration: ABe, DG, MES; Supervision: MLDM, DG, MES; Funding acquisition: DG, MES; Visualization: ABe; Writing – original draft: ABe, DG, MES; Writing – reviewing and editing: all authors.

## Supplemental material

### Supplementary figures

**Fig S1.**
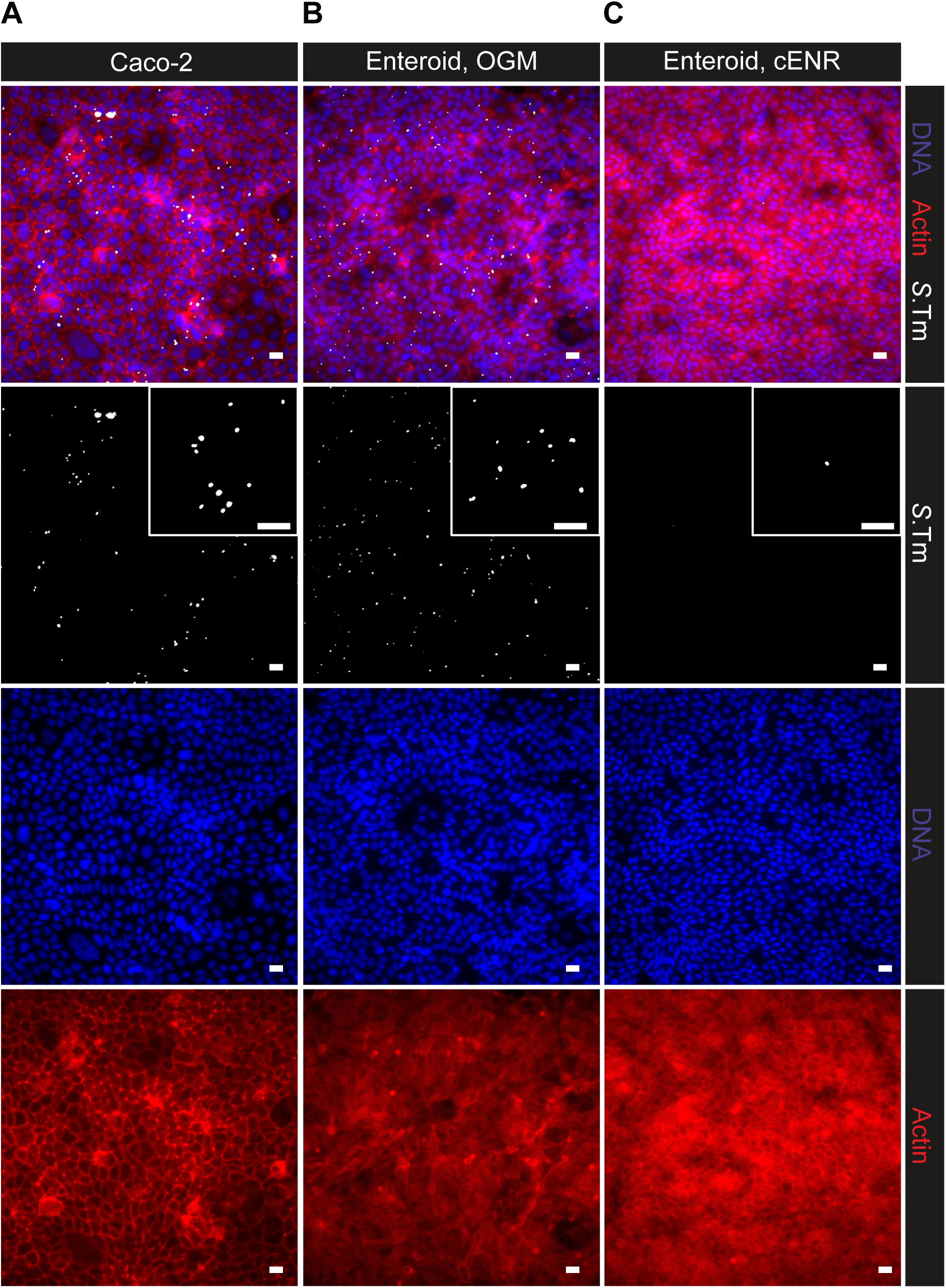
Fluorescence images from the platform for quantification of *S*.Tm epithelial cell invasion. Representative fluorescence images of IEC monolayers infected with *S*.Tm *wt* carrying the intracellular reporter construct p*ssaG*-GFP. **A.** Caco-2 cell monolayers, **B-C.** enteroid-derived monolayers maintained in **B.** organoid growth medium (OGM) and **C.** organoid differentiation medium (“cENR”; StemCell ODM). *S*.Tm foci are depicted in white. IECs were stained with DAPI (blue) and Alexa Fluor 647 Phalloidin (red). Scale bar: 20µm.

**Fig S2.**
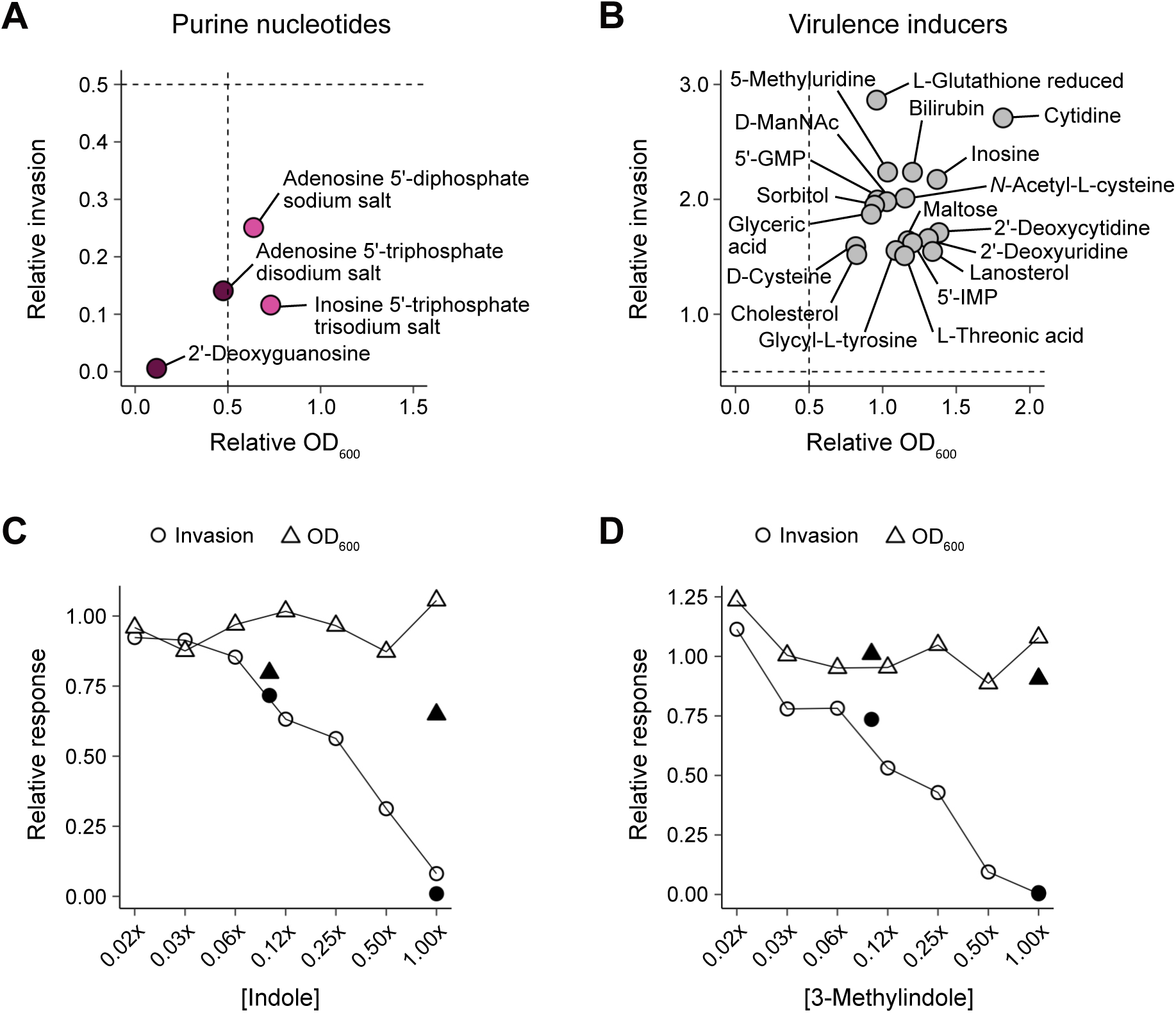
Screening of a gut metabolite library for anti-infective activities. **A-C.** *S*.Tm relative OD_600_ and relative invasion into IEC (Caco-2) monolayers after 4h pre-incubation with metabolites from the library: **A.** purine nucleotides and **B.** metabolites identified as virulence inducers (relative invasion >1.5). Dotted lines depict the 0.5 cut-offs of relative OD_600_ (x-axis) and relative invasion (y-axis). *N*-Acetyl-D-mannosamine (D-ManNAc), Guanosine 5’-monophosphate disodium salt (5’-GMP), Inosine-5’- monophosphate sodium salt hydrate (5’-IMP). **C-D.** Dose-response to **C.** indole and **D.** 3-methylindole starting at the highest concentration used in the main screen (“1x”). Black dots represent the corresponding results from the main metabolite library screen.

**Fig S3.**
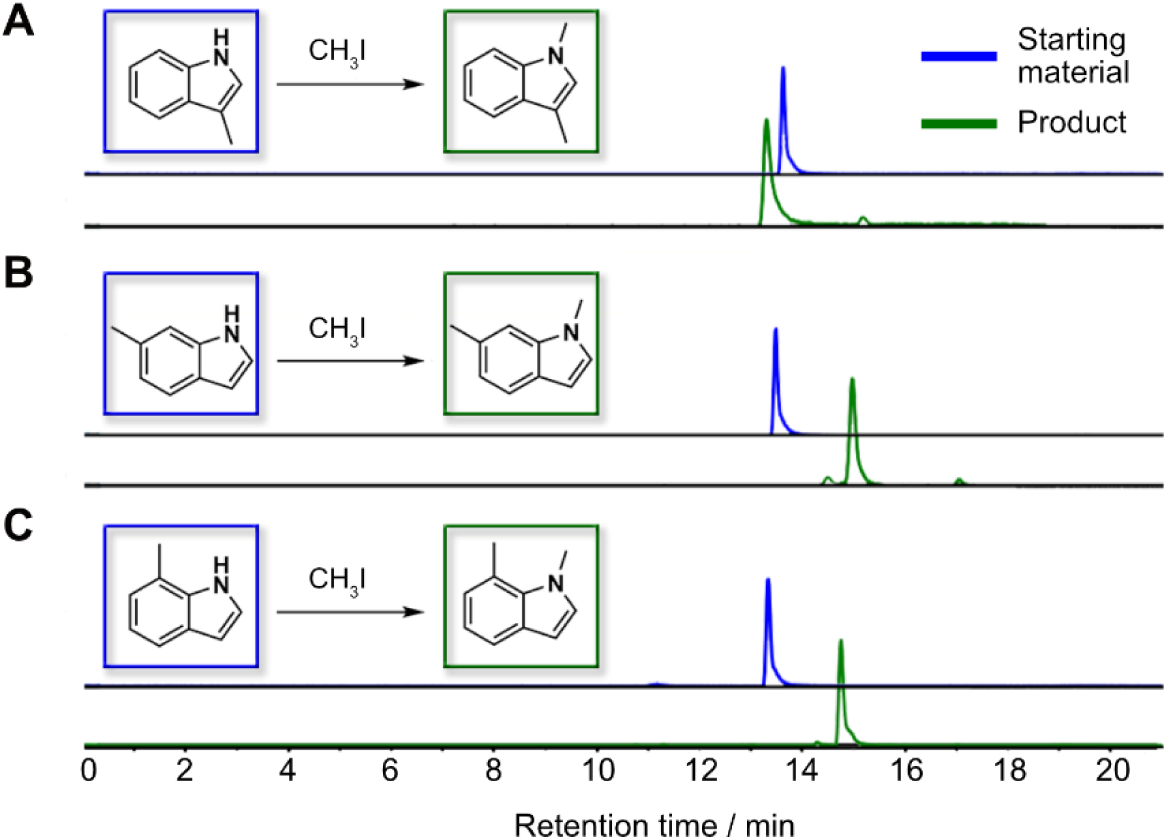
Synthesis of N-methylated indole derivatives. **A-C.** Illustration of synthesis of N-methylated indole-derivatives and retention time for starting material and product. **A.** 3-methylindole and 1,3-dimethylindole. **B.** 6-methylindole and 1,6-dimethylindole. **C.** 7-methylindole and 1,7-dimethylindole.

## Supplementary tables

**Table S1.** Strains and plasmids used in this study.

| Strain | Genotype | Reference |
| --- | --- | --- |
| *S.Tm wt* | SL1344, wt (SB300, Sm<sup>R</sup>) | (1) |
| *S.Tm ΔinvG* | SL1344, ΔinvG (SB161, Sm<sup>R</sup>) | (2) |
| *S.Tm ΔmotA* | SL1344, ΔmotA (SB300, Sm<sup>R</sup>) | (3) |
| *S.fl wt* | M90T, wt | (4) |
| *S.fl ΔmxiD* | M90T, ΔmxiD (Kan<sup>R</sup>) | (5) |
| Plasmid | Description | Reference |
| pM975 | pssaG-GFP (SPI2-dependent, vacuolar) | (6,7) |
| pZ1400 | puhpT-GFP (cytosolic) | (8,9) |

**Table S2.**
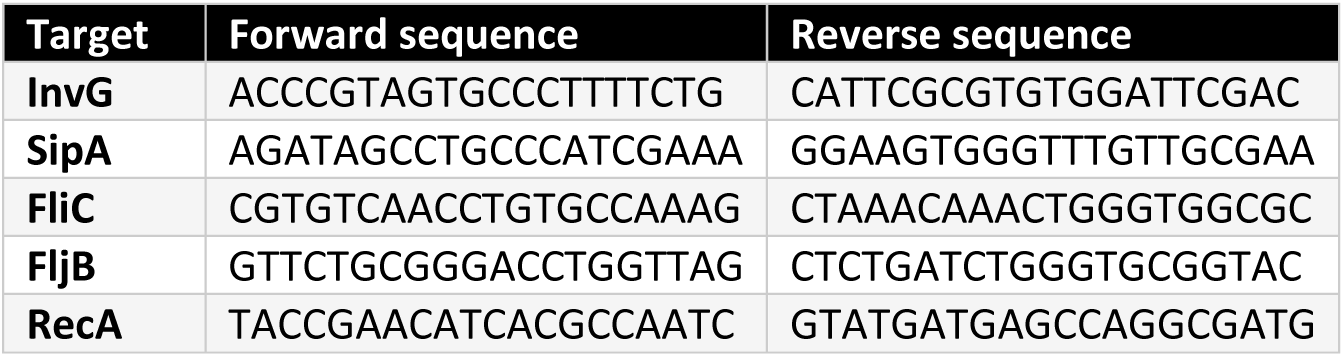
Primers used in this study.

### Supplementary chemical synthesis

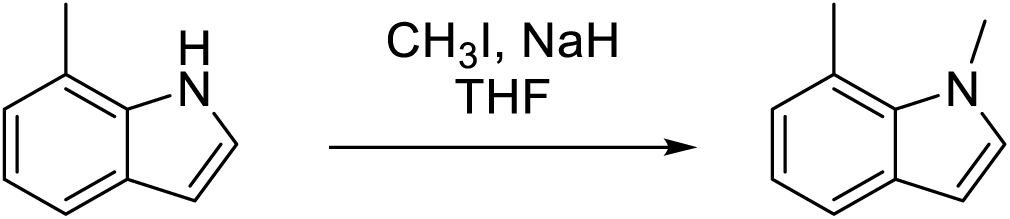

### Synthesis of 1,7-dimethylindole

Methylation of indole was done according to the literature procedure. To a solution of indole (50 mg) in THF (30 mL) at 0 °C was added NaH (13.7 mg, 60 % dispersion in mineral oil). The heterogeneous mixture was stirred at 0 °C for 15 minutes and 1h at room temperature. The mixture was then cooled to 0 °C, treated with iodomethane (31 μL), and allow to warm to room temperature. After 30 minutes, the reaction mixture was cooled to 0 °C, quenched with saturated NH4Cl (50 mL), and extracted with ether (3 x 30 mL). The organic layers were combined, washed with brine, dried over anhydrous Na2SO4, and concentrated in vacuo. The resulting oil was purified by flash chromatography (eluent: Hexane/EtOAc 10:1) to provide 1,7-dimethylindole (36.4 mg, 65.8%) as a colorless oil.

^1^H NMR (400 MHz, cdcl3) δ 7.49 (d, *J* = 7.8 Hz, 1H), 7.00 (t, *J* = 7.5 Hz, 1H), 6.96 (d, *J* = 3.1 Hz, 1H), 6.94 (d, *J* = 7.1 Hz, 1H), 6.47 (d, *J* = 3.1 Hz, 1H), 4.08 (s, 3H), 2.80 (s, 3H).

HRMS (ESI) *m/z* [M+H]⁺ calcd for C10H12N⁺ 146.0964; Found: 146.0965.

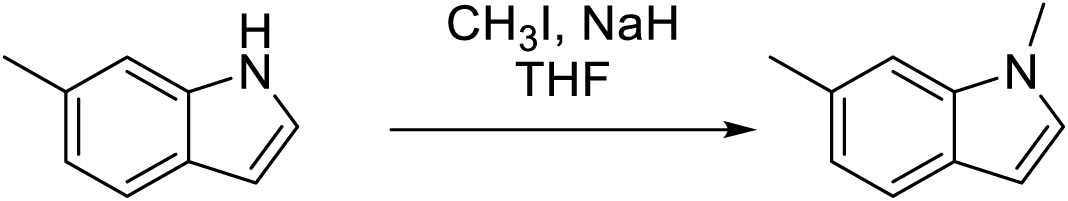

### Synthesis of 1,6-dimethylindole

Methylation of indole was done according to the literature procedure. To a solution of indole (50 mg) in THF (30 mL) at 0 °C was added NaH (13.7 mg, 60 % dispersion in mineral oil). The heterogeneous mixture was stirred at 0 °C for 15 minutes and 1h at room temperature. The mixture was then cooled to 0 °C, treated with iodomethane (31 μL), and allow to warm to room temperature. After 30 minutes, the reaction mixture was cooled to 0 °C, quenched with saturated NH4Cl (50 mL), and extracted with ether (3 x 30 mL). The organic layers were combined, washed with brine, dried over anhydrous Na2SO4, and concentrated in vacuo. The resulting oil was purified by flash chromatography (eluent: Hexane/EtOAc 10:1) to provide 1,6-dimethylindole (36.7 mg, 66.3%) as a colorless oil.

^1^H NMR (400 MHz, cdcl3) δ 7.56 (d, *J* = 8.0 Hz, 1H), 7.17 (s, 1H), 7.04 – 6.96 (m, 2H), 6.48 (d, *J* = 3.1 Hz, 1H), 3.78 (s, 3H), 2.55 (s, 3H).

HRMS (ESI) *m/z* [M+H]⁺ calcd for C10H12N⁺ 146.0964; Found: 146.0964.

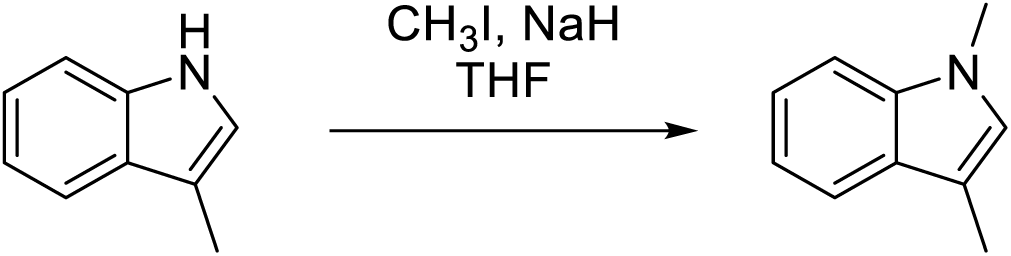

### Synthesis of 1,3-dimethylindole

Methylation of indole was done according to the literature procedure. To a solution of indole (50 mg) in THF (30 mL) at 0 °C was added NaH (13.7 mg, 60 % dispersion in mineral oil). The heterogeneous mixture was stirred at 0 °C for 15 minutes and 1h at room temperature. The mixture was then cooled to 0 °C, treated with iodomethane (31 μL), and allow to warm to room temperature. After 30 minutes, the reaction mixture was cooled to 0 °C, quenched with saturated NH4Cl (50 mL), and extracted with ether (3 x 30 mL). The organic layers were combined, washed with brine, dried over anhydrous Na2SO4, and concentrated in vacuo. The resulting oil was purified by flash chromatography (eluent: Hexane/EtOAc 10:1) to provide 1,3-dimethylindole (10.1 mg, 18.2%) as a colorless oil.

^1^H NMR (400 MHz, cdcl3) δ 7.56 (d, *J* = 7.9 Hz, 1H), 7.26 (t, *J* = 9.2 Hz, 1H), 7.21 (t, *J* = 7.5 Hz, 1H), 7.10 (t, *J* = 7.4 Hz, 1H), 6.81 (d, *J* = 1.3 Hz, 1H), 3.72 (s, 3H), 2.32 (s, 3H), 2.32 (d, *J* = 1.4 Hz, 3H).

HRMS (ESI) *m/z* [M+H]⁺ calcd for C10H12N⁺ 146.0964; Found: 146.0966.

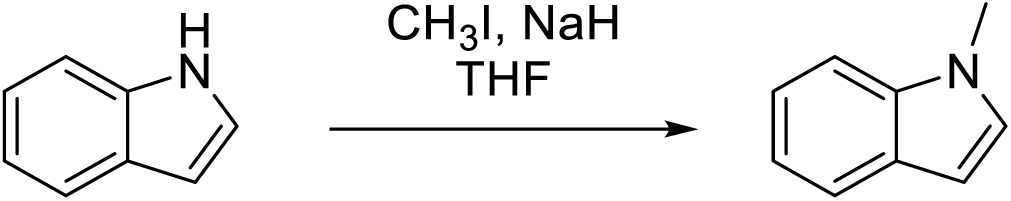

### Synthesis of 1-methylindole

Methylation of indole was done according to the literature procedure. To a solution of indole (50 mg) in THF (30 mL) at 0 °C was added NaH (13.7 mg, 60 % dispersion in mineral oil). The heterogeneous mixture was stirred at 0 °C for 15 minutes and 1h at room temperature. The mixture was then cooled to 0 °C, treated with iodomethane (31 μL), and allow to warm to room temperature. After 30 minutes, the reaction mixture was cooled to 0 °C, quenched with saturated NH4Cl (50 mL), and extracted with ether (3 x 30 mL). The organic layers were combined, washed with brine, dried over anhydrous Na2SO4, and concentrated in vacuo. The resulting oil was purified by flash chromatography (eluent: Hexane/EtOAc 10:1) to provide 1-methylindole (14.7 mg, 26.3%) as a colorless oil.

^1^H NMR (400 MHz, cdcl3) δ 7.65 (d, *J* = 7.8 Hz, 1H), 7.37 – 7.32 (m, 1H), 7.27 – 7.21 (m, 1H), 7.15 –7.10 (m, 1H), 7.07 (d, *J* = 3.1 Hz, 1H), 6.50 (dd, *J* = 3.1, 0.9 Hz, 1H), 3.81 (s, 3H).

HRMS (ESI) *m/z* [M+H]⁺ calcd for C9H10N⁺ 132.0808; Found: 132.0810.

## Supplementary NMR Spectra

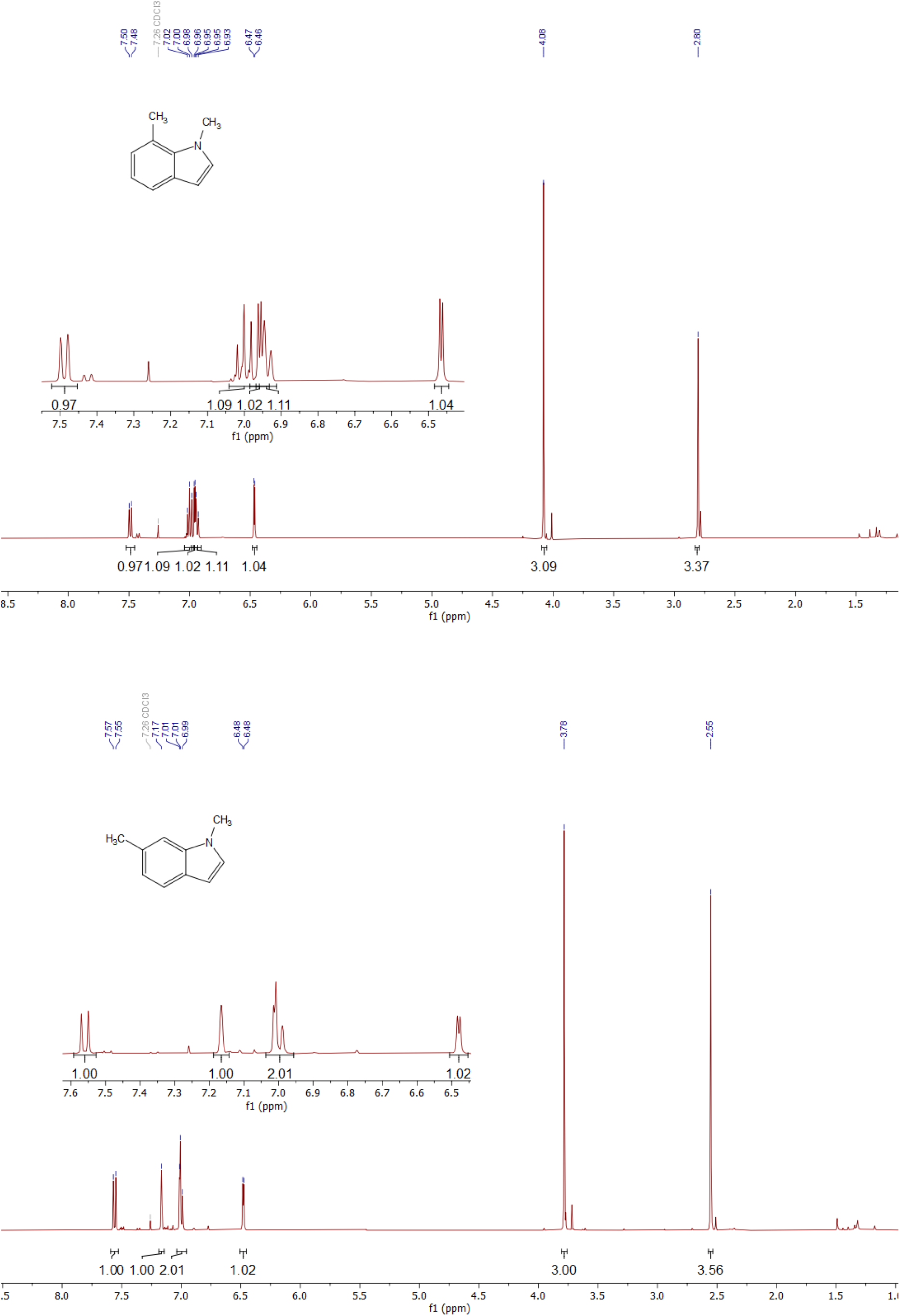
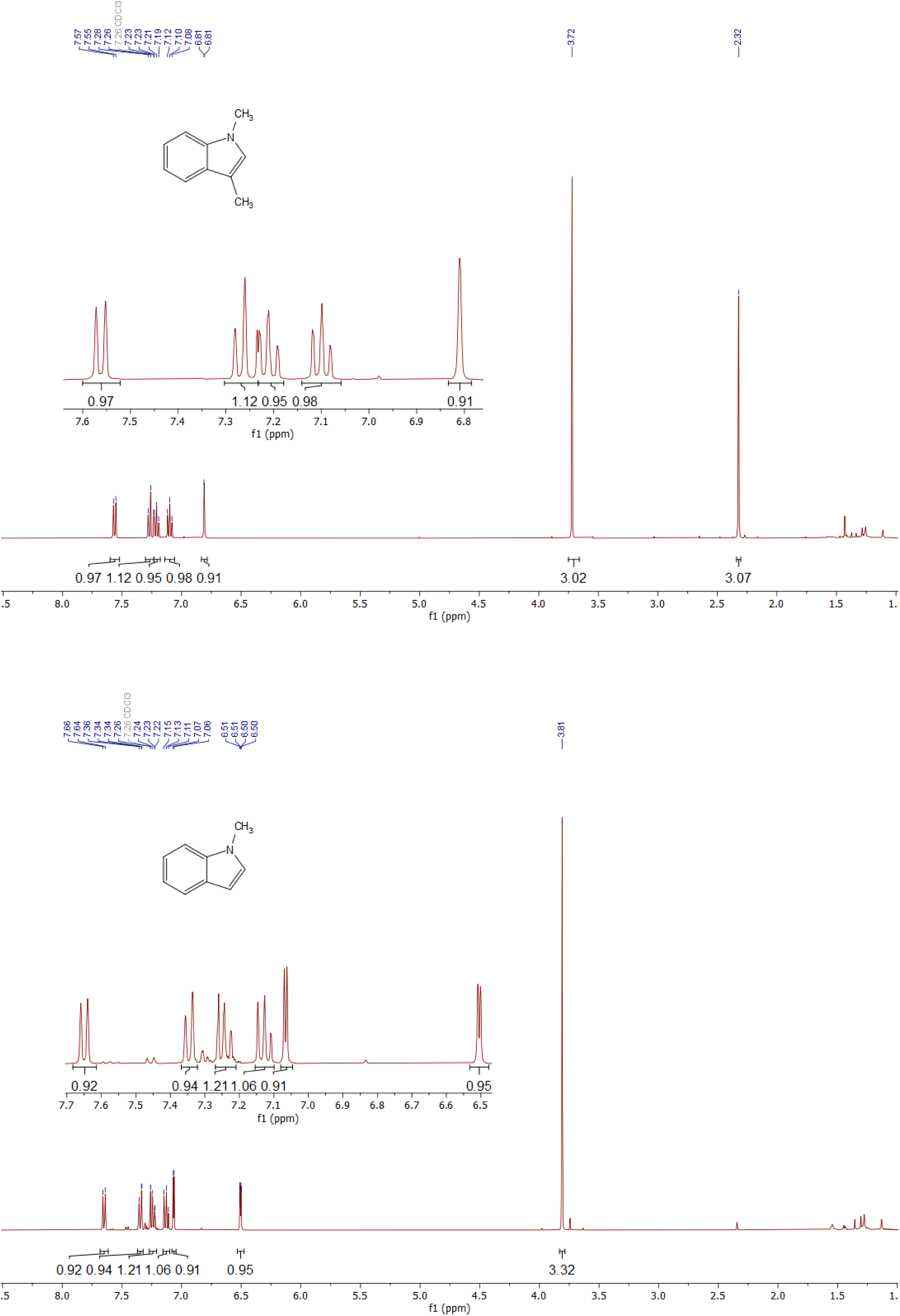

